# Continuous Motility Fingerprints from Gaussian Mixture Models Reveal Hidden Sperm Heterogeneity

**DOI:** 10.1101/2025.11.20.689493

**Authors:** Julianna Lamm

## Abstract

Computer-assisted sperm analysis (CASA) collapses sperm motility into coarse categorical bins, masking the heterogeneity that influences fertilization potential. Analyzing 33,500 trajectories from 340 individuals, we introduce a probabilistic fingerprinting framework that treats Gaussian mixture model (GMM) posteriors as soft supervision to construct continuous motility axes that capture forward progression, erratic movement, and uncertainty. This semi-supervised, posterior-guided approach condenses high-dimensional CASA features into interpretable, uncertainty-aware descriptors that capture both canonical and transitionary sperm behaviors. GMM consistently identified five reproducible subtypes, including an ‘erratic’ group characterized by vigorous, asymmetric motion associated with fertilization readiness. Patient-level fingerprints revealed robust associations with semen quality markers (debris, round cells, concentration), demonstrating potential utility for fertility evaluation and clinical decision-making. Together, our results provide the first large-scale evidence that probabilistic fingerprints capture biologically meaningful and clinically anchored sperm heterogeneity, establishing a generalizable framework for outcome-linked studies in reproductive medicine.

## 1. Introduction

Male sperm counts have dropped by 50% over the last 50 years (Levine et al., 2017), and male factors contribute to roughly half of infertility cases (NICHD, 2025). In the United States, demand for Assisted Reproductive Technology (ART) has more than doubled from 2013 to 2022 (CDC, 2024). There are two main ART procedures currently employed: in vitro fertilization (IVF),where tens of thousands of sperm are co-incubated with an oocyte, and intracytoplasmic sperm injection (ICSI), where a single sperm is selected and injected directly. Clinical practice has shifted toward ICSI, which directly bypasses sperm motility and quality barriers that limit fertilization in conventional IVF (Sciorio and Esteves, 2022).

Yet sperm are highly heterogeneous, and the challenge of assessing quality and selecting the most viable cells remains central to ART and fertility evaluation. Ejaculates contain diverse subpopulations with distinct motility profiles and current clinical assessments collapse this complexity into coarse aggregate percentages (Martínez-Pastor et al., 2011), obscuring variation that may influence fertilization success. Other fields have moved beyond such categorical scoring, for example, embryo grading now emphasizes continuous outcome-linked metrics (Papamentzelopoulou et al., 2024), and high-content assays like Cell Painting profile cells across continuous phenotypic axes (Bray et al., 2016).

Building on these precedents, we propose modeling sperm motility with Gaussian Mixture Models (GMM) and visualizing these profiles in continuous embedding spaces (e.g., UMAP, t-SNE). This approach moves beyond categorical CASA bins by preserving heterogeneity at both the sperm- and patient-levels. Soft posteriors enable the construction of granular “phenotypic fingerprints” that capture motility diversity within individuals and across cohorts. Such fingerprints provide a richer basis for downstream analysis and clinical translation. More broadly, posterior-based fingerprinting offers a general framework for linking single-cell heterogeneity to population-level health analytics.

## 2. Methods

### 2.1. Datasets

We analyzed 428 fully anonymized, de-identified sperm videos corresponding to 340 individuals, comprising 33,500 total tracks. At the track level, each sperm was represented by its (*x, y*) coordinates across frames and associated CASA kinematic metrics (VCL, VSL, VAP, LIN, STR, WOB, ALH, BCF, track length, and duration). At the patient level, an accompanying metadata file provided summary information for each video, including CASA grading percentages, number of views, debris count, round cell concentration, imaging resolution, sperm concentration, and semen volume. No patient identifiers were available. Tracks with fewer than *T*_min_ frames (default: 15) were excluded. Full definitions for CASA metrics are provided in Appendix A.1.

### 2.2. Clustering

We modeled sperm motility using Gaussian mixture models (GMM; primary) and k-means (baseline). Candidate values *k* ∈ 4, 5 were considered to align with canonical CASA categories (immotile, non-progressive, progressive, rapid-progressive) while allowing added granularity. With *k* = 5, a reproducible fifth cluster emerged with hyperactivated-like motion; Because true hyperactivation occurs only after capacitation (biochemical changes in the female reproductive tract that enable fertilization), we conservatively labeled this cluster “erratic” consistent with prior CASA cut-offs for hyperactivation (Mortimer and Swan, 1995).

For GMM, full covariance models were trained on MinMax-scaled CASA features, with model selection guided by BIC/AIC and qualitative interpretability. Each track *i* is represented by a posterior vector ***π***_*i*_ = [*P*_1_(*i*), …, *P*_*k*_(*i*)], which we treat as a continuous motility profile.

For k-means, clustering was performed on the same features with k-means++ initialization. Model selection relied on inertia (elbow) and silhouette diagnostics, and hard assignments were compared to GMM maximum-a-posteriori (MAP) labels using adjusted Rand index (ARI).

For visualization only, scaled features were embedded into 2D (UMAP, t-SNE) and colored by MAP labels.

### 2.3. Continuous motility axes

To move beyond categorical CASA bins, we derived three indices of sperm movement: *progressivity, erraticity*, and *entropy*. Inspired by Ben-Yehuda et al. (2022), who trained regressions on embryologist-annotated tracks, we instead used GMM posteriors as pseudo-labels in a semi-supervised framework. This approach collapses multivariate CASA features into smooth one-dimensional descriptors while preserving uncertainty. Full mathematical definitions are provided in Appendix A.3.

### 2.4. Validation and Robustness

We assessed robustness at the track, participant, and cohort levels using bootstrap and subsample resampling, and by projecting held-out tracks into the trained GMM space. Stability was quantified with adjusted Rand index (ARI) and cohort comparability with KL/Hellinger metrics, with full details in Appendix A.4. Analogous checks were performed for k-means.

## 3. Results

### 3.1. CASA bins obscure heterogeneity that GMM fingerprints recover

Across patients, GMM yielded a richer subtype distribution than CASA grading by recovering a substantial non-progressive population and separating *erratic* from *non-progressive* sperm (Figure 1). In contrast, CASA bins reported virtually no non-progressive sperm and conflated erratic trajectories with other categories, obscuring heterogeneity.

**Figure 1.**
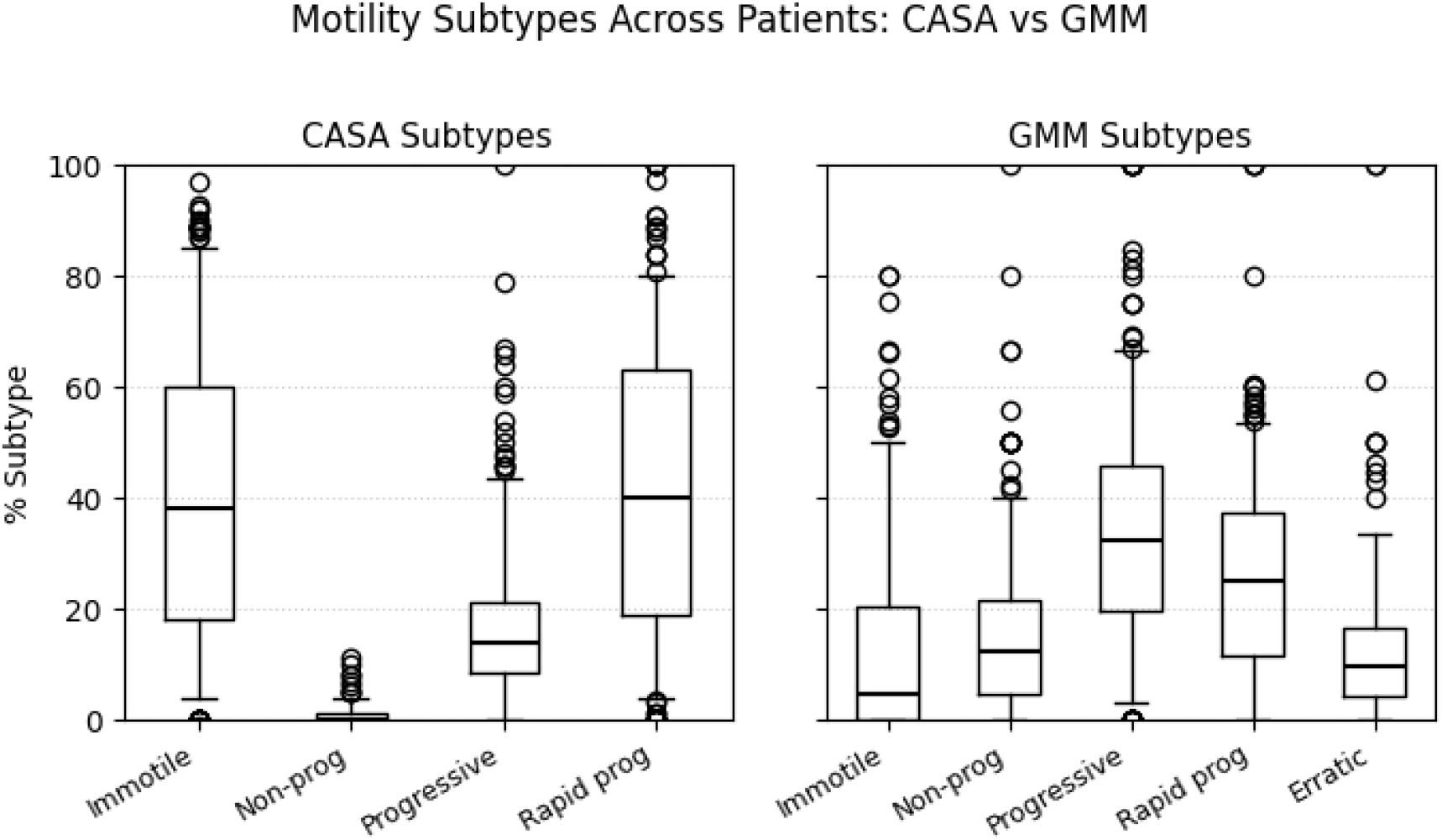
Motility subtype distributions across patients under CASA (left) and GMM (right). CASA bins obscure heterogeneity, reporting virtually no non-progressive sperm and conflating erratic trajectories, whereas GMM recovers both consistently.

These patterns were reproducible under bootstrap and participant-level resampling and reappeared in held-out projections, indicating they are not artifacts of the clustering procedure. Moreover, the recovered subtypes aligned with raw kinematic trends: non-progressive sperm showed slow, low forward progression, while erratic sperm exhibited excess lateral excursions—providing biological interpretability consistent with established motility behaviors (Table 1, Figure 2).

**Table 1:** Cluster centers from the *k* = 5 GMM solution, with assigned subtype labels. Corresponding k-means cluster centers are reported in Appendix Table B.2.

| ID | Label | ALH | BCF | LIN | MAD | STR | VAP | VCL | VSL |
| --- | --- | --- | --- | --- | --- | --- | --- | --- | --- |
| 0 | Progressive | 0.946 | 10.66 | 0.401 | 66.2 | 0.910 | 16.1 | 37.7 | 15.1 |
| 1 | Immotile | 0.264 | 11.69 | 0.217 | 46.2 | 0.724 | 2.7 | 10.1 | 2.0 |
| 2 | Erratic | 1.940 | 10.51 | 0.286 | 79.2 | 1.008 | 22.5 | 81.0 | 23.7 |
| 3 | Rapid progressive | 1.684 | 12.28 | 0.550 | 70.0 | 1.120 | 39.7 | 83.2 | 44.9 |
| 4 | Non-progressive | 1.092 | 14.09 | 0.675 | 63.3 | 1.100 | 39.5 | 64.2 | 43.1 |

**Figure 2.**
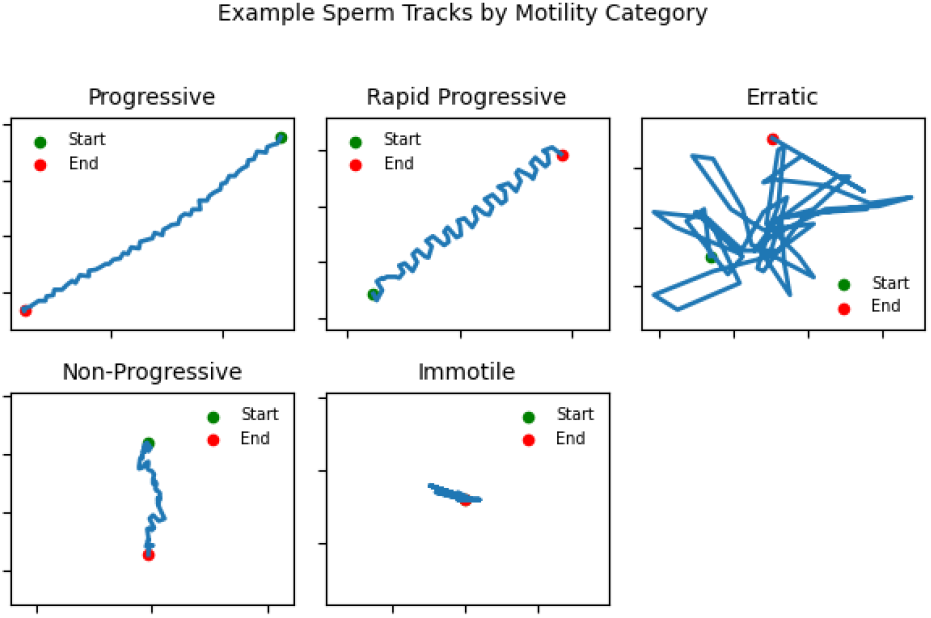
Representative sperm trajectories from the *k* = 5 GMM solution. The five clusters correspond to CASA’s four canonical motility categories plus an additional ‘erratic’ group characterized by rapid, asymmetric whiplike motion.

### 3.2. GMM recovers five reproducible motility subtypes

In addition to recovering heterogeneity overlooked by CASA, GMM consistently identified five reproducible motility subtypes across the cohort. These aligned with CASA’s four canonical categories (immotile, non-progressive, progressive, rapid progressive) while revealing a fifth “erratic” group characterized by rapid, asymmetric whip-like motion (Figure 2). This pattern resembles hyperactivated-like trajectories, but was not captured by CASA bins.

Across 50 bootstrap resamplings, GMM assignments were highly stable (median ARI = 0.96, IQR: 0.95–0.98), with similar robustness under participant-level resampling (median ARI = 0.91). Projection of held-out tracks into the trained embedding confirmed that cluster structures generalized to unseen samples. Although Silhouette values were low (see Appendix B.2), this outcome is expected given the biology of sperm motility. Individual sperm often exhibit mixed features of multiple movement types or transition between states within a single track, so their trajectories do not fit neatly into one category. This biological continuity reduces separation in standard clustering metrics, but posterior probabilities naturally capture such ambiguity by assigning partial membership and highlighting transitionary behavior. Cluster compositions corresponded closely to established CASA grading categories, with mean kinematic metrics for each cluster reported in Table 1. While k-means produced similar aggregate patterns, its reliance on hard assignments collapsed ambiguity into rigid bins. By contrast, GMM posteriors preserved heterogeneity and provided a foundation for constructing patient-level fingerprints. Detailed model selection curves, stability analyses, and generalization diagnostics are provided in Appendix B.1.

### 3.3. Semi-supervised motility axes capture clinically meaningful patterns

To move beyond rigid CASA bins, we developed continuous motility axes using a semi-supervised approach. GMM posteriors were treated as soft supervision, guiding linear regressions that collapsed CASA features into continuous axes of progressivity, erraticity, and entropy (Eqs. A.1–A.4). This posterior-guided construction collapses high-dimensional kinematic information into interpretable axes that preserve uncertainty and capture transitionary behaviors, providing clinically meaningful readouts of sperm motility.

#### 3.3.1. Single Cell Subtype Validation

The derived axes faithfully captured biologically interpretable motility patterns. The *progressivity index* increased in progressive (*ρ* = 0.51) and rapid progressive sperm (*ρ* = 0.38), while decreasing in immotile sperm (*ρ* = −0.64). The *erraticity index*, which weights excess ALH by posterior evidence, was maximized in the erratic subtype (*ρ* = 0.45) and negatively associated with progressive sperm (*ρ* = −0.53). The *entropy index*, quantifying posterior uncertainty, was highest in non-progressive sperm (*ρ* = 0.18), consistent with their heterogeneous, transitionary trajectories. Together, these results show that the semisupervised axes condense multivariate features into continuous descriptors that map cleanly onto known motility behaviors (Figure 3).

**Figure 3.**
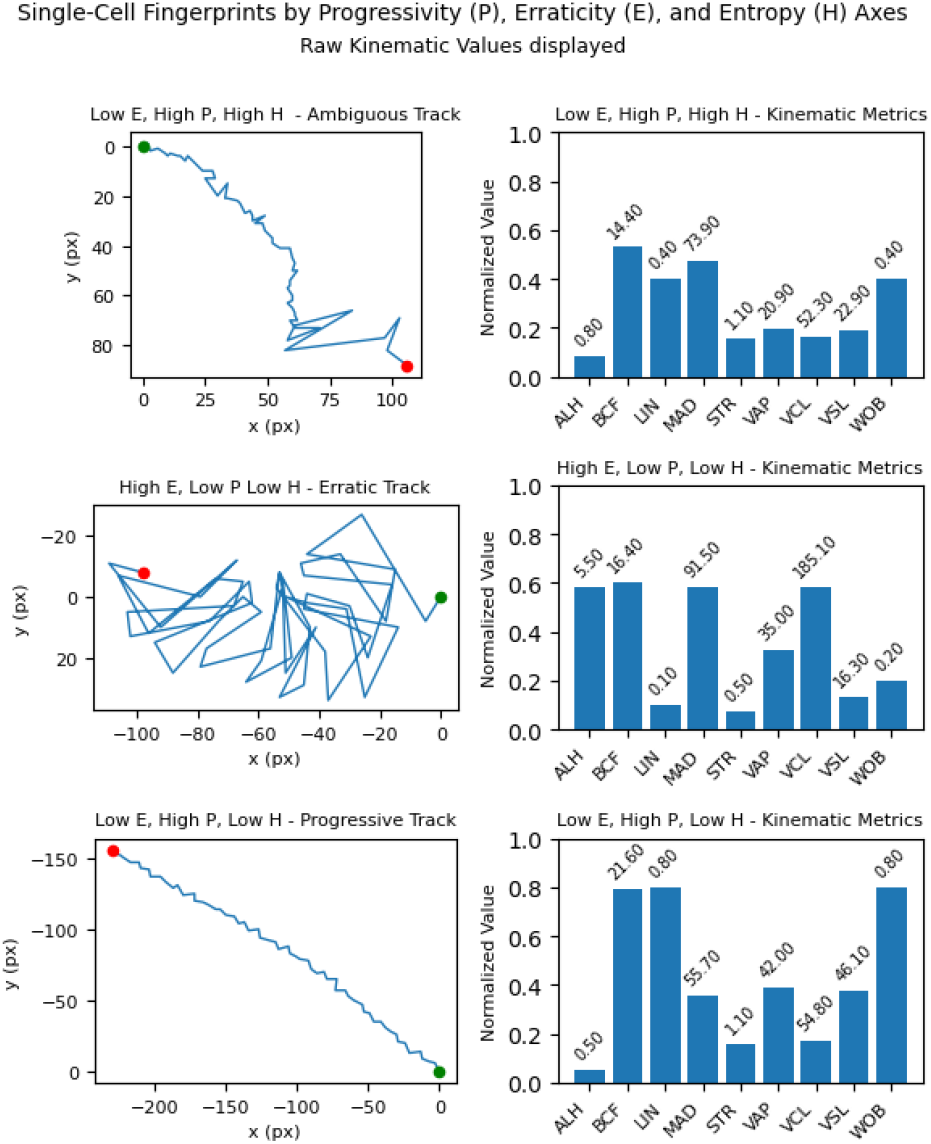
Representative single-cell fingerprints illustrating the semi-supervised motility indices. Left: sperm trajectories with corresponding progressivity (P), erraticity (E), and entropy (H) values. Right: associated kinematic feature profiles. Examples highlight (top) an ambiguous trajectory with high entropy, (middle) an erratic trajectory with high lateral excursions, and (bottom) a progressive trajectory with strong forward motion.

Plotting sperm in the progressivity–erraticity space, with entropy as color, highlighted both canonical motility categories and transitionary phenotypes (Figure 4). High-entropy cells populated intermediate regions, illustrating how posterior-guided axes capture uncertainty and continuous variation overlooked by categorical bins.

**Figure 4.**
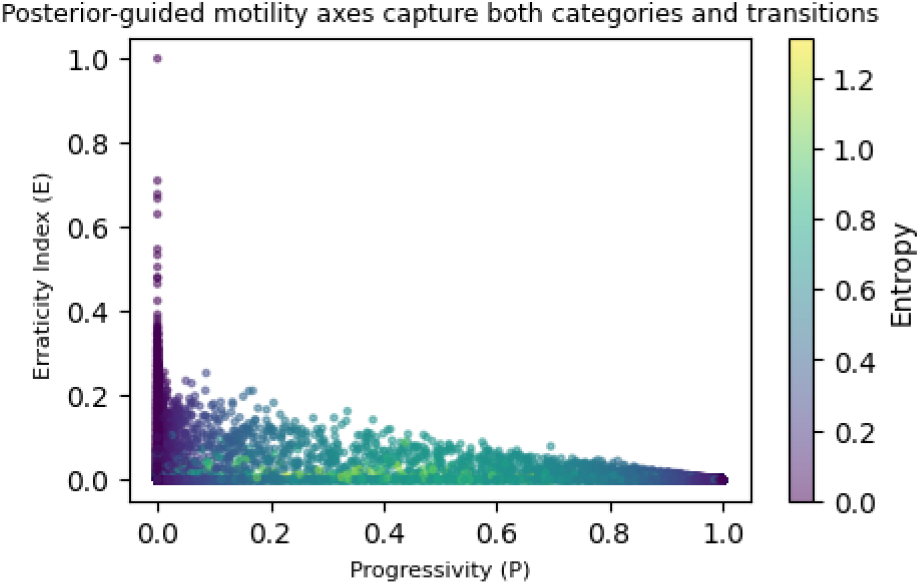
Posterior-guided motility axes at the single-cell level. Each point represents a sperm cell positioned by progressivity (P) and erraticity (E), colored by entropy (H). Canonical categories appear at the extremes, while high-entropy cells occupy intermediate regions, highlighting transitionary phenotypes.

#### 3.3.2. Composite fingerprints reveal patient-level motility archetypes

Having validated the axes at the single-cell level, we next aggregated them across patients to generate composite “fingerprints” of motility behavior. Each fingerprint integrates GMM subtype distributions, posterior entropy, mean CASA-derived metrics, and representative trajectories into a compact visualization (Figure 5). This multi-view representation enables direct comparison of individuals in terms of both overall motility balance and within-sample heterogeneity, revealing archetypes ranging from clean progressive profiles to heterogeneous mixtures with high uncertainty.

**Figure 5.**
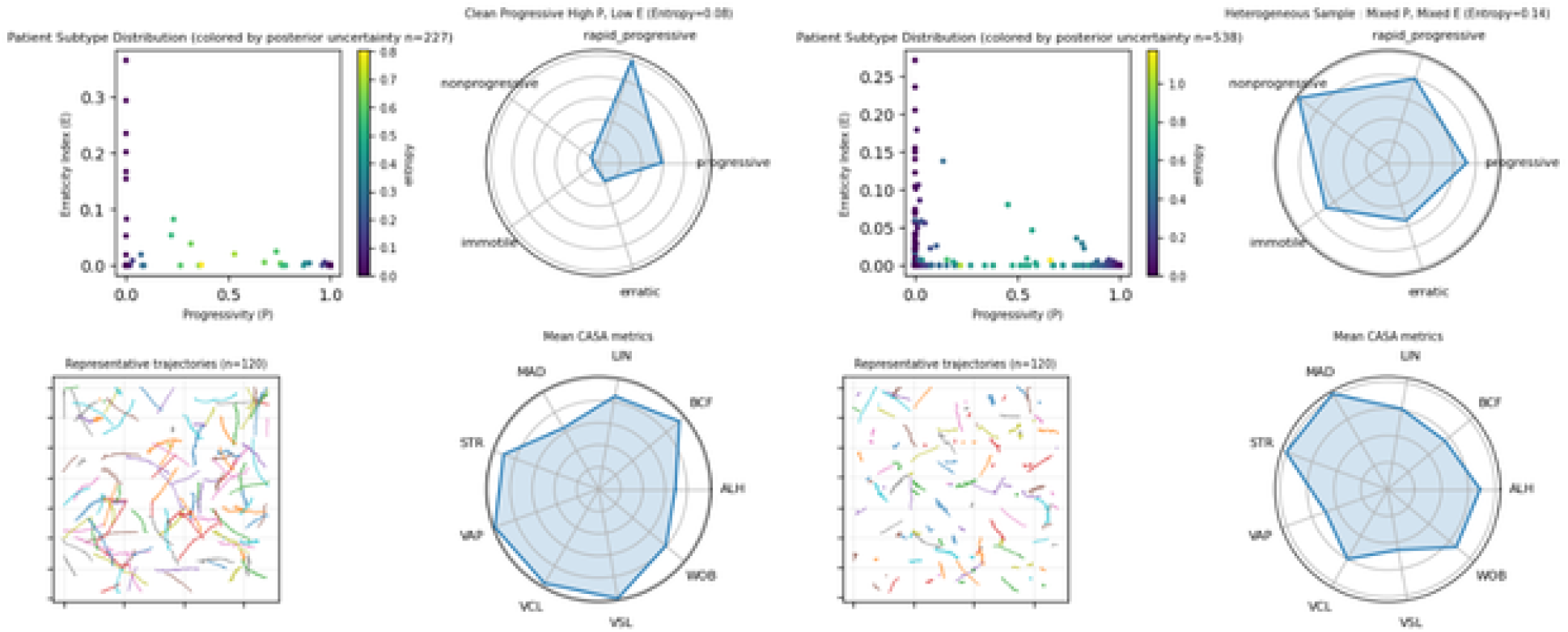
Composite patient-level motility fingerprints. Each panel summarizes an individual’s motility profile by integrating GMM subtype distributions, mean CASA-derived metrics (radar plots), representative trajectories, and posterior entropy. This compact visualization highlights between-patient differences in overall motility balance as well as within-sample heterogeneity, revealing archetypes ranging from clean progressive profiles to heterogeneous mixed patterns with high uncertainty.

**Figure 6.**
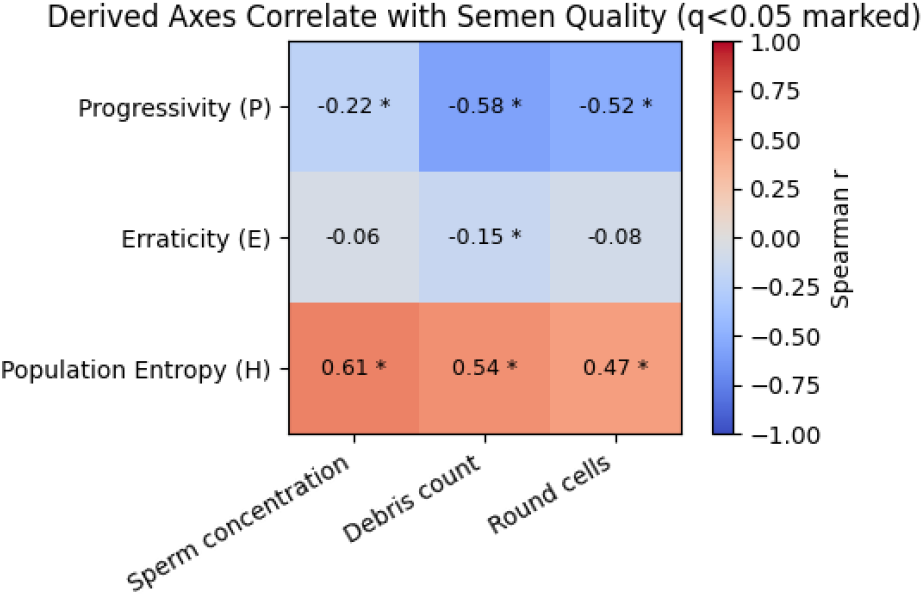
Spearman correlations between derived motility indices (progressivity, erraticity, entropy) and semen quality markers. Significant associations (*q* < 0.05) are marked, with progressivity negatively and entropy positively correlated with contamination and concentration.

**Figure 7.**
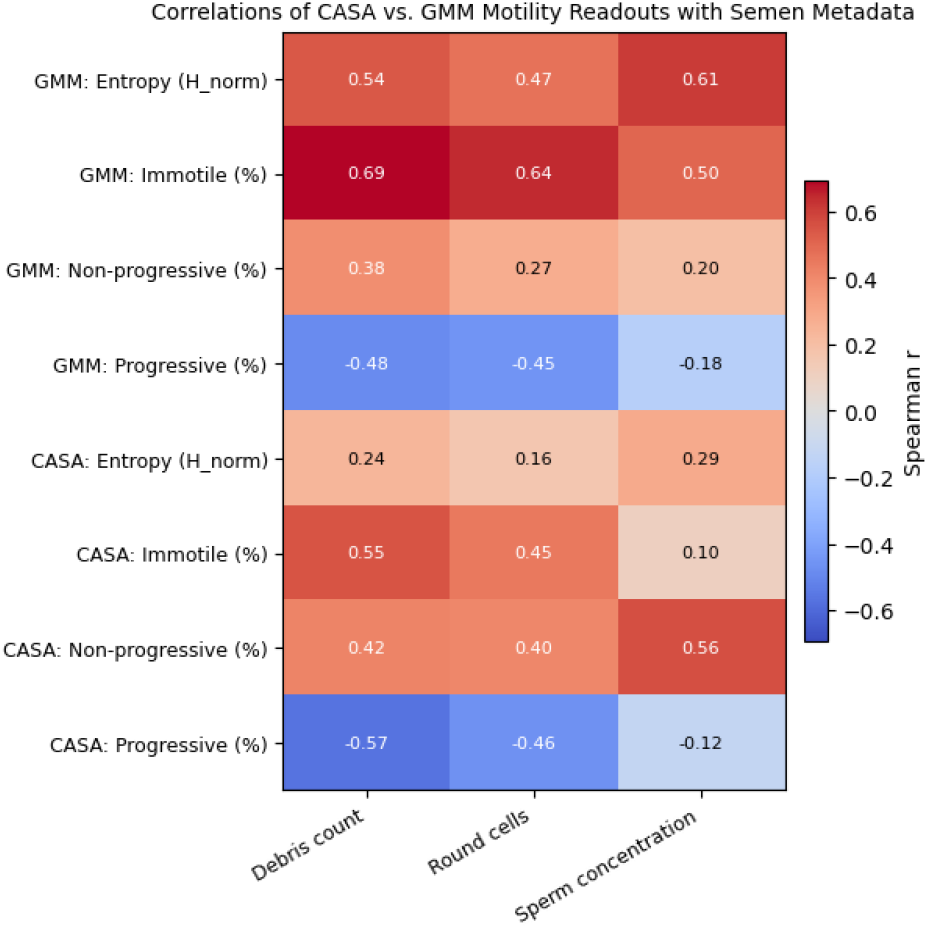
Spearman correlations between semen quality metadata (debris, round cells, sperm concentration) and motility readouts. GMM-derived features show stronger and more consistent associations than CASA percentages.

#### 3.3.3. Metadata Correlations

To assess clinical relevance, we tested whether patient-level fingerprints correlated with semen metadata (sperm concentration, debris count, and round cell concentration). GMM-derived features showed stronger and more consistent associations than CASA (Figure 3.3.3). The *progressivity index* was negatively correlated with debris (*ρ* = −0.58, *q* < 0.05) and round cells (*ρ* = −0.52, *q* < 0.05), consistent with impaired forward motion in contaminated samples. In contrast, *population entropy* was positively correlated with all three measures (*ρ* = 0.47–0.61, *q* < 0.05), reflecting greater motility heterogeneity in lower-quality semen. Percent immotile also tracked strongly with debris and round cells (*ρ* = 0.64–0.69, *q* < 0.05).

For comparison, we examined CASA-derived features (Figure 3.3.3). While CASA immotility also correlated with debris, GMM-derived entropy and subtype balances provided stronger and more consistent anchoring to metadata, suggesting that our fingerprints capture clinically meaningful dimensions not well reflected in CASA cutoffs.

Together, these results indicate that continuous axes and GMM fingerprints not only capture interpretable motility variation but also track with independent clinical indicators of semen quality more effectively than CASA.

## 4. Discussion

Our analysis provides the first large-scale evidence that sperm motility heterogeneity can be reproducibly captured using probabilistic fingerprints rather than categorical CASA bins. Across 33,500 tracks, Gaussian mixture models consistently recovered a fifth “erratic” subtype resembling hyperactivated-like motion, alongside clinically interpretable continuous axes of progressivity, erraticity, and entropy. To the best of our knowledge, the ability to reproducibly extract this movement pattern has not been demonstrated at scale and holds particular promise for identifying sperm with enhanced fertilization potential. These readouts not only aligned with raw kinematic trends but also tracked robustly with semen quality markers, underscoring their biological validity and translational potential.

While outcome linkage and capacitation ground truth remain open directions, our results demonstrate that heterogeneity overlooked by CASA is stable, quantifiable, and clinically anchored. By establishing reproducible motility subtypes and clinically relevant continuous axes, this study lays a foundation for integrating motility diversity into future reproductive assessments and outcome-linked research. More broadly, our findings highlight how uncertainty-aware machine learning can reveal meaningful biological variation in health data that conventional scoring schemes obscure.

Several limitations warrant mention, including the absence of capacitation ground truth, reliance on standard CASA features without explicit site-effect modeling, and lack of direct outcome linkage. Nonetheless, posterior-weighted summaries and entropy provide richer readouts of sample quality and variability than rigid CASA bins. In sum, posterior fingerprints of sperm motility recover biologically meaningful subtypes, collapse them into continuous descriptors, and align with metadata of clinical relevance.

## Data and Code Availability

This study used anonymized, de-identified sperm videos and derived features that are not publicly available and code is not released.

## Institutional Review Board (IRB)

This work involved secondary analysis of anonymized, deidentified sperm videos and features provided for research. No identifiers were available, and we were not involved in data collection. Accordingly, this does not constitute human subjects research and did not require IRB approval.

## Appendix A. Additional Methods

### A.1. CASA kinematic metrics

The following standard CASA kinematic metrics were used:

- **VCL**:Curvilinear velocity: speed along the actual head trajectory (µm/s).
- **VSL**: Straight-line velocity: displacement from start to end divided by time (µm/s).
- **VAP**: Average path velocity: speed along a smoothed average trajectory (µm/s).
- **LIN**: Linearity: VSL/VCL (dimensionless).
- **STR**: Straightness: VSL/VAP (dimensionless).
- **WOB**: Wobble: VAP/VCL (dimensionless).
- **ALH**: Amplitude of lateral head displacement: mean side-to-side excursion (µm).
- **BCF**: Beat-cross frequency: number of crossings of the average path per second (Hz).
- **MAD**: Mean angular displacement: average change in trajectory angle (deg).
- **Track length**: Number of frames per track.
- **Duration**: Time span of the track (s).

Units assume pixel–to–µm calibration and video frame rate for conversion.

#### A.2. CASA grading percentages

Standard CASA grading percentages as defined by World Health Organization (WHO) were provided as metadata for each patient:

- **Grade A (Rapid progressive)** Spermatozoa moving forward quickly in a straight line with a velocity of at least 25 µm/s at 37°C.
- **Grade B (Slow or sluggish progressive)** Spermatozoa with forward movement that is either slower, less straight, or both. Their velocity is typically between 5 and 25 µm/s.
- **Grade C (Non-progressive)** Motile sperm that exhibit movement but fail to move forward. This includes sperm that swim in tight circles or whose flagella only beat with no forward progression.
- **Grade D (Immotile)** Spermatozoa that show no movement.

Reported values are percentages of total counted sperm falling into each category.

**Table B.1:** Clustering and embedding hyperparameters.

| Method | Settings |
| --- | --- |
| GMM | Full cov.; $k=4, 5$; BIC/AIC |
| K-means | k-means++; 50 restarts |
| UMAP | $n_{\text{nbrs}}=15$; $\min_{\text{dist}}=0.1$ |
| t-SNE | Perplexity = 30 |

#### A.3. Mathematical definitions of continuous motility axes

##### A.3.1. Progressivity axis

We defined a soft ordinal target by assigning increasing weights to GMM posteriors:

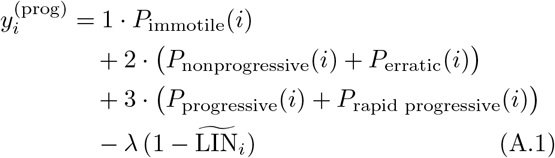

A linear regression was then trained on z-scored CASA features to predict 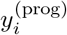. Model outputs were normalized to [0, 1].

##### A.3.2. Erraticity axis (regression-based)

Erraticity was defined as:

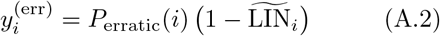

A second linear regression was trained on z-scored CASA features to predict 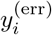, with outputs normalized to [0, 1].

##### A.3.3. Erraticity (residual-based check)

As a complementary feature-engineering check, we modeled expected ALH as a function of linearity (LIN) and straight-line velocity (VSL). Residuals were computed as:

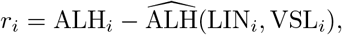

where 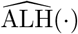 is the fitted baseline. Residuals were z-scored, rectified (negative values set to zero), and weighted by the erratic posterior:

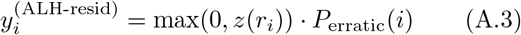

This metric highlights excess lateral excursions beyond expected values, serving as a simple, interpretable validation of the regression-based erraticity axis.

##### A.3.4. Per-Sperm Posterior Entropy (Shannon Entropy)

Per-sperm entropy quantified uncertainty of GMM posterior assignments:

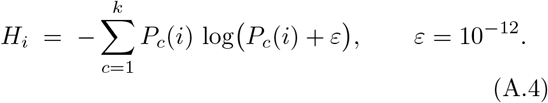

Low entropy indicates confident subtype assignments, while high entropy reflects ambiguous motility phenotypes. At the patient level, entropy was aggregated across sperm to provide a summary measure of heterogeneity.

##### A.3.5. Patient-level Entropy (Mean Shannon Entropy)

Entropy was aggregated across all tracks for patient *p* to provide a measure of sample heterogeneity:

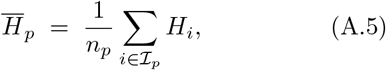

where *ℐ*_*p*_ indexes the sperm tracks for patient *p* and *n*_*p*_ = | *ℐ*_*p*_|.

#### A.4. Validation and robustness

##### A.4.1. Model selection

For a dataset *X* ∈ ℝ^*n×d*^, features were first scaled to [0, 1] using a MinMax transform. For each candidate number of clusters *k* ∈ {*k*_min_, …, *k*_max_}, a Gaussian mixture model (GMM) was fit and scored using the Akaike information criterion (AIC) and Bayesian information criterion (BIC):

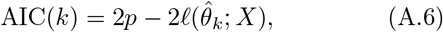

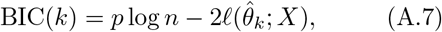

where *p* is the number of free parameters and 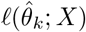 is the maximized log-likelihood. Model choice was guided by arg min_*k*_ BIC(*k*) together with an “elbow” criterion, defined as the first *k* for which the relative BIC improvement fell below a threshold *τ* :

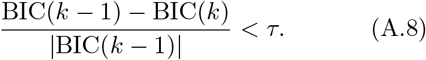

##### A.4.2. Track-level resampling

Baseline MAP labels 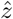 were obtained by fitting a GMM to the full dataset. For each bootstrap iteration *b* = 1, …, *B*, a resample *X*^(*b*)^ of size *m* = ⌈*αn*⌉ was drawn (with or without replacement). A new GMM was fit to *X*^(*b*)^, and labels 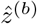 were predicted for the full dataset. Stability was quantified using the adjusted Rand index (ARI):

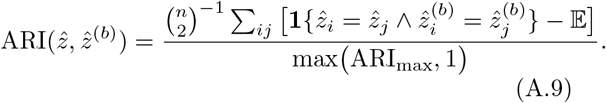

##### A.4.3. Participant-level resampling

Let 𝒫 denote the set of all participants, with |𝒫| total. At each iteration, we randomly sampled a subset of approximately *ρ* |𝒫| participants, where *ρ* ∈ (0, 1) is the sampling fraction (here *ρ* = 0.7). Since the number of participants must be an integer, this was rounded down:

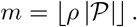

A GMM was then fit using only tracks from the selected participants, and predicted labels on the full dataset were compared to baseline assignments using ARI (Eq. A.9).

##### A.4.4. Generalization to held-out data

For a held-out set *X*_test_, posterior distributions *π*^held-out^ were computed using the scaler and GMM trained on *X*_primary_. Let 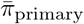 and 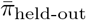 denote the mean subtype composition vectors. Cohort comparability was assessed using symmetric KL divergence and Hellinger distance:

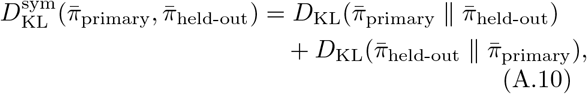

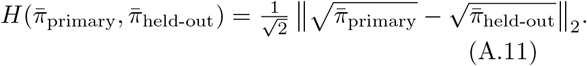

## Appendix B. Supplementary Results and Figures

### B.1. Model Selection and Validation Metrics

#### B.2. K-means Cluster Centers

**Figure B.1:**
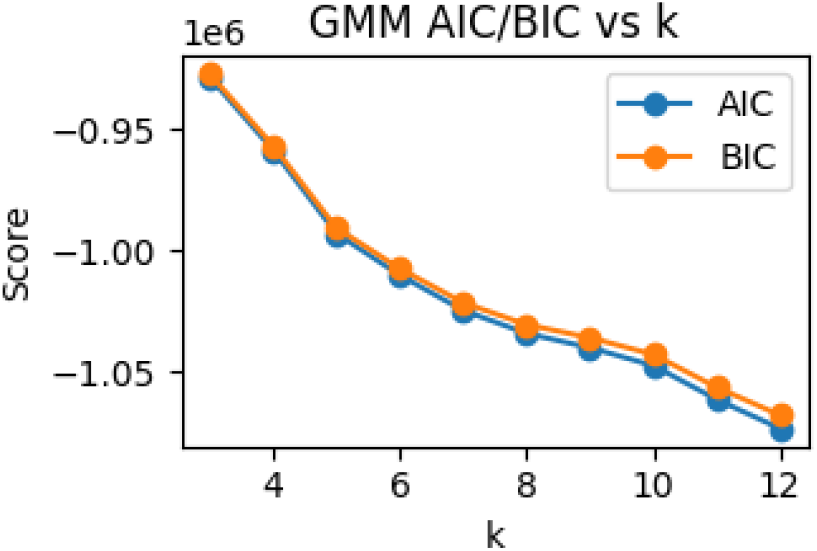
GMM model selection. AIC and BIC values across candidate cluster numbers *k*. Both criteria decrease monotonically without a clear cutoff.

**Figure B.2:**
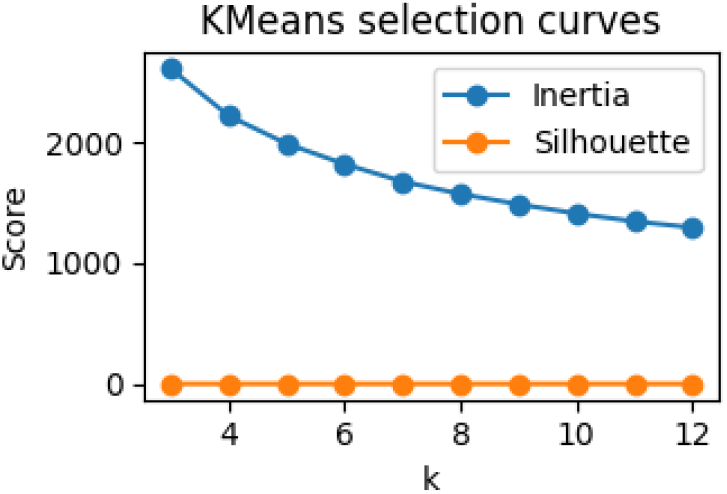
K-means model selection. Inertia decreases monotonically with *k*, while silhouette scores remain uniformly low, offering limited guidance for cutoff selection.

**Table B.2:** Internal clustering indices (Silhouette, Davies-Bouldin (DB), Calininski-Harabasz (CH).

| Split | Silhouette | CH | DBI |
| --- | --- | --- | --- |
| Primary | 0.091 | 8071.22 | 2.170 |
| Held-out | 0.075 | 588.66 | 2.182 |

**Figure B.3:**
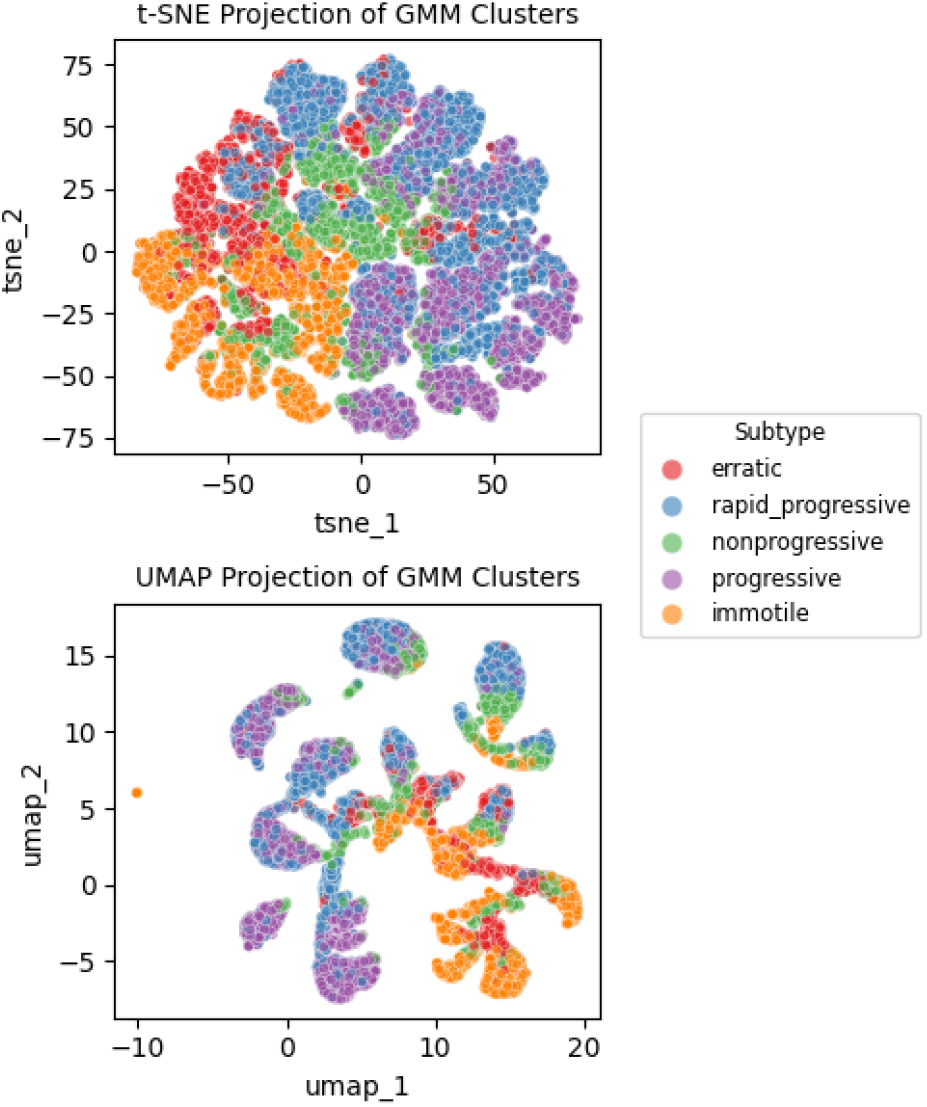
Two-dimensional embeddings of sperm tracks using t-SNE (top) and UMAP (bottom), colored by GMM cluster assignment.Clusters show broad separation with expected overlap, reflecting the continuous spectrum of sperm motility phenotypes.

**Figure B.4:**
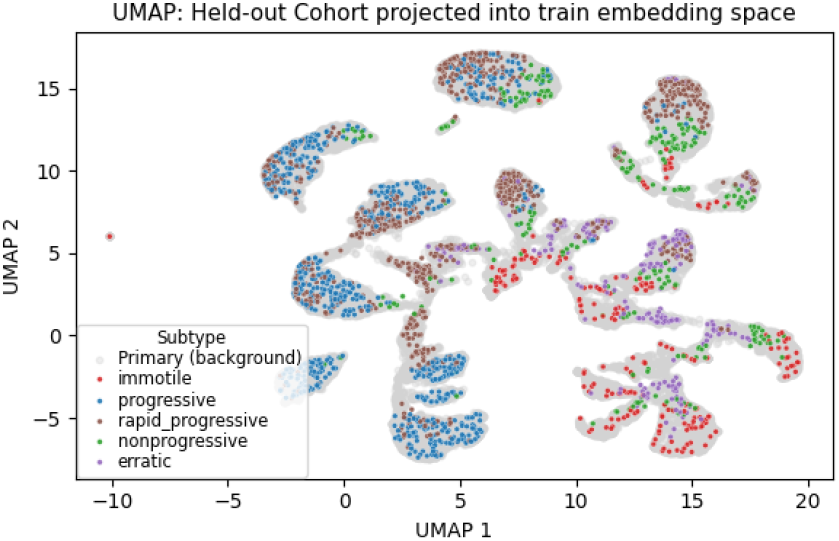
Projection of held-out tracks into the UMAP embedding learned on primary data. Cluster structure is preserved, indicating generalization beyond the initial dataset.

**Table B.3:** K-means cluster centers (part 1): ALH, BCF, LIN, MAD, STR.

| Subtype | ALH | BCF | LIN | MAD | STR |
| --- | --- | --- | --- | --- | --- |
| Progressive | 0.979 | 15.60 | 0.773 | 56.02 | 1.144 |
| Rapid prog | 1.759 | 12.79 | 0.544 | 74.47 | 1.135 |
| Non-prog | 0.881 | 10.37 | 0.605 | 55.05 | 1.131 |
| Immotile | 0.467 | 11.64 | 0.173 | 53.57 | 0.658 |
| Erratic | 1.889 | 10.86 | 0.323 | 83.08 | 0.980 |

**Table B.4:**
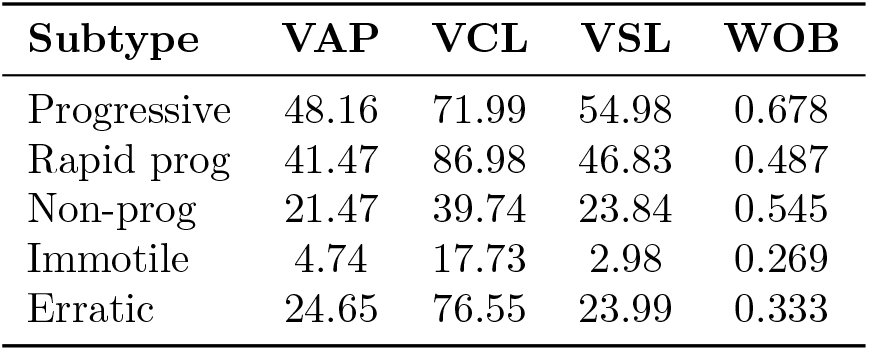
K-means cluster centers (part 2): VAP, VCL, VSL, WOB.

## References

K. Ben-Yehuda, S. K. Mirsky, M. Levi, I. Barnea, I. Meshulach, S. Kontente, D. Benvaish, R. Cur-Cycowicz, Y. N. Nygate, and N. T. Shaked. Simultaneous morphology, motility, and fragmentation analysis of live individual sperm cells for male fertility evaluation. Advanced Intelligent Systems, 4: 2100200, 2022. doi: 10.1002/aisy.202100200.

M. A. Bray, S. Singh, H. Han, et al. Cell painting, a high-content image-based assay for morphological profiling using multiplexed fluorescent dyes. Nature Protocols, 11: 1757–1774, 2016. doi: 10.1038/nprot.2016.105.

CDC. National assisted reproductive technology (art) summary. https://www.cdc.gov/art/php/national-summary/, 2024. Centers for Disease Control and Prevention; accessed 2025-09-09.

Hagai Levine, Niels Jørgensen, Alexandre Martino-Andrade, Jaime Mendiola, Dana Weksler-Derri, Inbar Mindlis, Roberta Pinotti, and Shanna H Swan. Temporal trends in sperm count: a systematic review and meta-regression analysis. Human Reproduction Update, 23(6): 646–659, 2017. doi: 10.1093/humupd/dmx022.

Felipe Martínez-Pastor, E. Jorge Tizado, J. Julian Garde, Luis Anel, and Paulino de Paz. Statistical series: Opportunities and challenges of sperm motility subpopulation analysis. riogenology, 75(5): 783–795, 2011. ISSN 0093-691X. doi: 10.1016/j.theriogenology.2010.11.034. URL https://www.sciencedirect.com/science/article/pii/S0093691X10006278.

S. T. Mortimer and M. A. Swan. Kinematics of capacitating human spermatozoa analysed at 60 hz. Human Reproduction, 10(4):873–879, April 1995. doi: 10.1093/oxfordjournals.humrep.a136053.

NICHD. How common is male infertility, and what are its causes? https://www.nichd.nih.gov/health/topics/menshealth/conditioninfo/infertility, 2025. Eunice Kennedy Shriver National Institute of Child Health and Human Development. Accessed: 2025-09-09.

M. S. Papamentzelopoulou, I. N. Prifti, D. Mavrogianni, T. Tseva, N. Soyhan, A. Athanasiou, A. Athanasiou, A. Athanasiou, P. Vogiatzi, G. Konomos, D. Loutradis, and M. Sakellariou. Assessment of artificial intelligence model and manual morphokinetic annotation system as embryo grading methods for successful live birth prediction: A retrospective monocentric study. Reproductive Biology and Endocrinology, 22(1):27, March 2024. doi: 10.1186/s12958-024-01198-7.

R. Sciorio and S. C. Esteves. Contemporary use of icsi and epigenetic risks to future generations. Journal of Clinical Medicine, 11(8):2135, April 2022. doi: 10.3390/jcm11082135.

